# Relationship between luteal size and plasma progesterone concentration in pregnant and non-pregnant Indian crossbred dairy cows

**DOI:** 10.1101/2023.07.02.547387

**Authors:** Sanjay Agarwal, Harihar Prasad Gupta, Shiv Prasad, Irfan Ahmad Khan, Afroza Khanam, Firdous Ahmad Khan

**Affiliations:** Department of Veterinary Gynaecology and Obstetrics, College of Veterinary and Animal Science, G. B. Pant University of Agriculture and Technology, Pantnagar, India; Division of Veterinary Gynaecology and Obstetrics, Faculty of Veterinary Science and Animal Husbandry, R. S. Pura, India; Department of Large Animal Medicine and Surgery, School of Veterinary Medicine, St. George’s University, True Blue, Grenada

**Author notes:** <u>**Correspondence:**</u> Firdous A. Khan, BVSc, MVSc, DVSc, Diplomate ACT.

**Keywords:** Bovine, corpus luteum, ultrasonography, pregnancy, progesterone

## Abstract

The relationship between luteal size and plasma progesterone concentration in purebred cows has been investigated previously. However, there is limited information on this topic in Indian crossbred cows. This study aimed to evaluate the relationship between luteal size parameters (diameter and area) and plasma progesterone concentration in healthy pregnant and non-pregnant Indian crossbred cows. Fifty healthy lactating crossbred cows were artificially inseminated after estrus detection and retrospectively classified into pregnant (n=29) and non-pregnant (n=21) based on the results of ultrasonographic pregnancy diagnosis. Data on luteal diameter, luteal area, and plasma progesterone concentration collected on Days 7, 14, and 20 in non-pregnant cows and on Days 7, 14, 20, 25, 30, 35, 40, and 60 in pregnant cows were analyzed. Spearman’s correlation test was used to evaluate the association between the corpus luteum size parameters and the plasma progesterone concentration. Luteal area was positively correlated with plasma progesterone concentration in both pregnant (ρ=0.48, P<0.001) and non-pregnant (ρ=0.43, P=0.001) cows. Luteal diameter had a relatively weaker correlation with plasma progesterone concentration in both pregnant (ρ=0.45, P<0.001) and non-pregnant (ρ=0.24, P=0.058) cows. These results suggest that luteal area is a better indicator of CL function than luteal diameter in healthy Indian crossbred cows.

## Introduction

Corpus luteum (CL) plays a pivotal role in the regulation of the estrous cycle and maintenance of pregnancy in cattle. Therefore, it is not surprising that luteal size and circulating progesterone concentration are frequently studied endpoints in bovine reproduction research. Luteal size in cattle is measured using transrectal ultrasonography (Pierson *et al*., 1988) and commonly expressed in terms of either luteal diameter (Taylor and Rajamahendran, 1991; Ricci *et al*., 2017; Zwiefelhofer *et al*., 2020) or luteal area (Kastelic *et al*., 1990; Ginther *et al*., 2012; Pugliesi *et al*., 2012; Rocha *et al*., 2019). Circulating progesterone concentrations are commonly determined using enzyme-linked immunosorbent assay (ELISA) or radioimmunoassay (RIA) of plasma or serum harvested from peripheral blood samples (Ginther *et al*., 2012; Pugliesi *et al*., 2012; Broes and LeBlanc, 2014). Several studies on the association between luteal size and circulating progesterone have been reported in pregnant and non-pregnant purebred cattle (Kastelic *et al*., 1990; Siqueira *et al*., 2009; de Tarso *et al*., 2016; Berger *et al*., 2017; Vrisman *et al*., 2018; Rocha *et al*., 2019). Similar information is lacking in Indian crossbred cattle that constitute about 26.6% of the total cattle population in in the country as per the latest census report published by the Department of Animal Husbandry and Dairying in 2019. Therefore, the objective of this study was to evaluate the relationship between luteal size parameters (diameter and area) and plasma progesterone concentration in healthy pregnant and non-pregnant Indian crossbred cows.

## Materials and methods

The Institutional Ethics Committee (IAEC/VGO/CVASc/181) and the Committee for the Purpose of Control and Supervision of Experiments on Animals (CPCSEA) approved all animal handling protocols and experimental procedures used in this study.

### Animals and breeding management

Fifty healthy cyclic Indian crossbred dairy cows (4 to 7 years old, at least 60 days in milk, BCS 3 to 4) were enrolled. The enrolled cows were selected from a research herd after conducting transrectal ultrasonography and the white side test in order to only include cows that did not have any apparent reproductive abnormalities. Daily visual observation for behavioral signs of standing estrus was conducted in the morning and evening. Transrectal ultrasonography was performed to examine cows that were suspected to be in estrus. Cows with a preovulatory follicle (≥ 10 mm diameter) were bred twice through AI at 12 and 24 hrs after standing estrus. Pregnancy diagnosis was conducted using ultrasonography and the cows were retrospectively classified as pregnant (n=29) or non-pregnant (n=21). All cows were maintained under uniform husbandry conditions (housing and feeding) during the study.

### Corpus luteum size and plasma progesterone concentration

Corpus luteum diameter and area were measured using built-in measurement functions in a portable ultrasound scanner (DIGI-600 M, PROVET; S.S. Medical Systems (I) Private Limited, India) equipped with a 5.0 MHz linear-array transducer. Progesterone concentration was measured using plasma samples harvested from blood collected in heparinized test tubes through jugular venipuncture. Commercial radioimmunoassay kits (M/s Beckman Coulter IM 1188) with analytical sensitivity of 0.03 ng/mL and including a highly specific progesterone antibody were used for progesterone estimation. Data on luteal diameter, luteal area, and plasma progesterone concentration collected on Days 7, 14, and 20 in non-pregnant cows and on Days 7, 14, 20, 25, 30, 35, 40, and 60 in pregnant cows were used to evaluate the association between corpus luteum size and plasma progesterone concentration.

## Statistical analysis

Commercial statistical software (SPSS Version 16.0, SPSS Inc., Chicago, IL, USA) was used to conduct statistical analysis of the data. The data were tested for normality through the Shapiro-Wilk test, the results of which indicated that the data were not normally distributed. The relationships between corpus luteum size parameters and plasma progesterone concentration were evaluated using Spearman’s correlation test. Statistical significance was set at P-values less than 0.05. Results are presented as Spearman’s correlation coefficients and their associated P-values.

## Results and Discussion

For each of the three parameters (luteal diameter, luteal area, and plasma progesterone concentration), a total of 63 observations in non-pregnant cows and 232 observations in pregnant cows were analyzed. There was a significant positive correlation between luteal area and plasma progesterone concentration in non-pregnant cows. In contrast, luteal diameter had a weak positive correlation with plasma progesterone concentration that approached significance. Luteal diameter and luteal area were positively correlated with each other, and the correlation was highly significant (Table 1). In pregnant cows, relatively stronger positive correlations were observed between luteal diameter, luteal area, and plasma progesterone concentration (Table 2). Similar to the observation in non-pregnant cows, luteal area was slightly more strongly correlated with plasma progesterone concentration than luteal diameter.

**TABLE 1.**
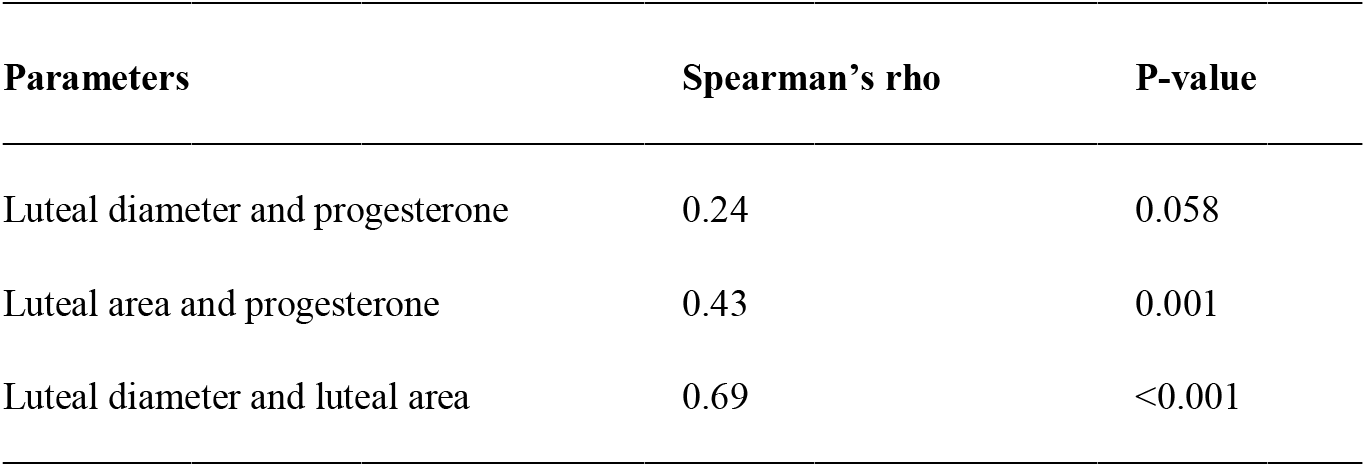
Correlations between luteal diameter, luteal area, and plasma progesterone concentration in non-pregnant Indian crossbred cows (n=21)

**TABLE 2.** Correlations between luteal diameter, luteal area, and plasma progesterone concentration in pregnant Indian crossbred cows (n=29)

| Parameters | Spearman's rho | P-value |
| --- | --- | --- |
| Luteal diameter and progesterone | 0.45 | <0.001 |
| Luteal area and progesterone | 0.48 | <0.001 |
| Luteal diameter and luteal area | 0.88 | <0.001 |

The associations between luteal size parameters and plasma progesterone concentration observed in the Indian crossbred cows in this study are similar to those reported previously in purebred dairy and beef cattle (Kastelic *et al*., 1990; Siqueira *et al*., 2009; de Tarso *et al*., 2016; Berger *et al*., 2017; Vrisman *et al*., 2018; Rocha *et al*., 2019). Our results suggest that luteal area may be a better indicator of CL function than luteal diameter in pregnant and non-pregnant Indian crossbred cows. Future studies including luteal size parameters and blood flow are required to further our understanding of the relationship between luteal structure, vascularity, and function in Indian crossbred cows.

## Data availability statement

The data that support the findings of this study are available from the corresponding author upon reasonable request.

## Acknowledgements

The authors would like to acknowledge the assistance provided by the staff of the IDF, Nagla, GBPUAT, Pantnagar, India with handling and restraint of the cows during this study.

## Author contributions

Sanjay Agarwal: Conceptualization, Methodology, Investigation. Harihar Prasad Gupta: Funding Acquisition, Project Administration, Supervision. Shiv Prasad: Project Administration, Resources, Supervision. Irfan Ahmad Khan: Writing – Review and Editing. Afroza Khanam: Data Curation, Visualization, Writing – Original Draft. Firdous Ahmad Khan: Conceptualization, Formal analysis, Software, Writing – Review and Editing.

## References

Berger, H, Lietzau, M, Tichy, A and Herzog, K (2017). Pregnancy outcome is influenced by luteal area during diestrus before successful insemination but not by milk production level. Theriogenology. 104: 115–119.

Broes, A and LeBlanc, SJ (2014). Comparison of commercial progesterone assays for evaluation of luteal status in dairy cows. Can. Vet. J., 55: 582–584.

De Tarso, SGS, Apgar, GA, Gastal, MO and Gastal, EL (2016). Relationships between follicle and corpus luteum diameter, blood flow, and progesterone production in beef cows and heifers: preliminary results. Anim. Reprod., 13: 81–92.

Ginther, OJ, Khan, FA, Hannan, M.A, Rodriguez, MB, Pugliesi, G and Beg, MA (2012). Role of LH in luteolysis and growth of the ovulatory follicle and estradiol regulation of LH secretion in heifers. Theriogenology. 77: 1442–1452.

Kastelic, JP, Bergfelt, DR and Ginther, OJ (1990). Relationship between ultrasonic assessment of the corpus luteum and plasma progesterone concentration in heifers. Theriogenology. 33: 1269–1278.

Pierson, RA, Kastelic, JP and Ginther, OJ (1988). Basic principles and techniques for transrectal ultrasonography in cattle and horses. Theriogenology. 29: 3–20.

Pugliesi, G, Khan, FA, Hannan, MA, Beg, MA, Carvalho, GR and Ginther, OJ (2012). Inhibition of prostaglandin biosynthesis during post luteolysis and effects on CL regression, prolactin, and ovulation in heifers. Theriogenology. 78: 443–454.

Ricci, A, Carvalho, PD, Amundson, MC and Fricke, PM (2017). Characterization of luteal dynamics in lactating Holstein cows for 32 days after synchronization of ovulation and timed artificial insemination. J. Dairy Sci., 100: 9851–9860.

Rocha, CC, Martins, T, Cardoso, BO, Silva, LA, Binelli, M and Pugliesi, G (2019). Ultrasonography-accessed luteal size endpoint that most closely associates with circulating progesterone during the estrous cycle and early pregnancy in beef cows. Anim. Reprod. Sci., 201: 12–21.

Siqueira, LG, Torres, CA, Amorim, LS, Souza, ED, Camargo, LS, Fernandes, CA and Viana, JH (2009). Interrelationships among morphology, echotexture, and function of the bovine corpus luteum during the estrous cycle. Anim. Reprod. Sci., 115: 18–28.

Taylor, C and Rajamahendran, R (1991). Follicular dynamics, corpus luteum growth and regression in lactating dairy cattle. Can. J. Anim. Sci., 71: 61–68.

Vrisman, DP, Bastos, NM, Rossi, GF, Rodrigues, NN, Borges, L, Taira, AR, de Paz, C, Nogueira, GP, Teixeira, P, Monteiro, FM and Oliveira, M (2018). Corpus luteum dynamics after ovulation induction with or without previous exposure to progesterone in prepubertal Nellore heifers. Theriogenology. 106: 60–68.

Zwiefelhofer, EM, Davis, BM and Adams, GP (2020). Research and development of a silicone letrozole-releasing device to control reproduction in cattle. Theriogenology. 146: 104–110.

